# Differential TM4SF5-mediated SIRT1 modulation and signaling for chronic liver disease

**DOI:** 10.1101/2020.02.19.956151

**Authors:** Jihye Ryu, Eunmi Kim, Min-Kyung Kang, Dae-Geun Song, Eun-Ae Shin, Jae Woo Jung, Seo Hee Nam, Ji Eon Kim, Hye-Jin Kim, Jeong-Hoon Lee, Jung-Hwan Yoon, Taekwon Son, Semi Kim, Hwi Young Kim, Jung Weon Lee

## Abstract

Here we show the roles of transmembrane 4 L six family member 5 (TM4SF5) in the progression of nonalcoholic steatosis (or NAFL) to steatohepatitis (NASH). The overexpression of TM4SF5 caused nonalcoholic steatosis and NASH in an age-dependent manner. Initially, TM4SF5-positive hepatocytes and livers exhibited lipid accumulation, decreased SIRT1, increased SREBPs levels, and inactive STAT3 via SOCS1/3 upregulation. In older animals, TM4SF5 under an inflammatory environment increased SIRT1 expression and STAT3 activity with no significant change to SOCSs and SREBPs levels, leading to active STAT3-mediated fibrotic extracellular matrix (ECM) production. Liver tissues from clinical human patients with NAFL or NASH also showed such a TM4SF5-SIRT1-STAT3-ECM relationship correlated with fibrosis score and age. Ligand-independent and TM4SF5-mediated STAT3 activity led to collagen I and laminins/laminin γ2 expression in hepatic stellate cells and hepatocytes, respectively. Laminin γ2 suppression abolished CCl_4_-mediated liver damage and ECM production and reduced SIRT1 and active-STAT3, but did not alter SREBP1 or SOCSs levels. These findings suggest that TM4SF5, CCL20, SIRT1, and/or laminin γ2 may be promising therapeutic targets against liver disease.

## Introduction

Chronic liver injury causes inflammation and metabolic dysregulation, leading to the progression of liver diseases(Tu et al, 2015). Nonalcoholic liver injury occurs due to immune-mediated or direct injury to hepatocytes, resulting first in hepatic fat deposition (steatosis, non-alcoholic fatty liver; NAFL), then in inflammation (steatohepatitis), and finally in excessive extracellular matrix (ECM) deposition(Gong et al, 2017). Excessive ECM deposition results in abnormal cell proliferation and scar tissue (fibrosis/cirrhosis) and eventually leads to hepatocellular carcinoma (HCC)(Bonnans et al, 2014).

Current disease models of nonalcoholic steatohepatitis (NASH) reveal excess metabolic dysfunction in the liver, resulting in cell stress, inflammation, and fibrosis(Sanyal, 2019). The metabolic roles of hepatocytes cannot thus be ignored when investigating the liver pathogenesis. The immune-metabolic aspects of nonalcoholic fatty liver disease (NAFLD) and NASH can be investigated as approaches to block the progression to HCC(Anstee et al, 2019; Sircana et al, 2019). Nonalcoholic injury-mediated liver inflammation causes production of diverse cytokines(Biancheri et al, 2014) and chemokines(Marra & Tacke, 2014), which may facilitate ECM production and subsequent fibrosis. CCL2 (MCP-1), CCL5 (RANTES), CCL20 (MIP-3α), and CXCL10 (IP-10) are involved in NAFLD/NASH and fibrosis(Marra & Tacke, 2014). Hepatic ECM depositions are produced primarily by hepatic stellate cells (HSCs)(Gressner & Weiskirchen, 2006), although hepatocytes, which compose 85% of the liver mass(Roskams T, 2007), also cause ECM depositions(Tu et al, 2015). Fibrous septa is composed mainly of collagen I depositions produced by activated HSCs(Maher & McGuire, 1990). However, *in vitro* hepatocytes also produce ECMs. Therefore, it is reasonable to investigate the roles of other hepatic cell and ECM types in liver fibrosis(Tu et al, 2015).

Transmembrane 4 L six family member 5 (TM4SF5) is an *N-*glycosylated membrane protein with four transmembrane domains(Lee, 2015) that is induced by TGFβ1 signaling during CCl_4_-mediated liver fibrosis(Kang et al, 2012b) and highly expressed in liver cancer(Lee et al, 2008). TM4SF5 activates c-SRC via a direct physical association(Jung et al, 2013), and thereby activates STAT3(Ryu et al, 2014). Hepatic TM4SF5 further modulates mTOR and S6K1 activity, both of which are critical for cell proliferation(Jung et al, 2019). Thus, TM4SF5 can activate intracellular signaling molecules during liver disease progression from fibrosis/cirrhosis to HCC. To understand if TM4SF5 could also be involved in the progression from NAFL to NASH would thus facilitate the discovery of treatments that can prevent NASH itself and further the progression to hepatic fibrosis/cirrhosis or HCC.

Here, the roles of TM4SF5 in NAFLD were explored using transgenic, knockout, and diet- or chemical-induced mouse models and their primary cells. We found that TM4SF5-transgenic mice developed NAFL, NASH, and fibrotic phenotypes in an age-dependent manner. We hypothesized that TM4SF5-mediated signaling pathways supported progression of liver malignancy. We characterized here how the signaling axes of TM4SF5-mediated SIRT1 modulation toward either SREBPs or SOCSs/STAT3 regulated lipid and ECM metabolism, respectively, to promote progression from steatosis to NASH and fibrosis.

## Results

### TM4SF5 overexpression caused nonalcoholic fatty liver disease (NAFLD)

Unlike wild-type (WT) mice, systemic TM4SF5-overexpressing transgenic mice (C57BL/6-Tg^TM4SF5^) exhibited abdominal obesity, fatty liver (Fig. 1A), and greater lipid accumulation in the liver (Fig. 1B) at one year of age. Furthermore, Tg^TM4SF5^ mouse livers exhibited elevated levels of hepatic triacylglycerol (TG) and alanine aminotransferase (GPT/ALT) compared with WT livers; however, albumin levels were unchanged (Fig. 1C). Thus, TM4SF5 overexpression caused phenotypes resembling NAFLD (Fig. 1D). mRNA and protein levels of acetyl-CoA carboxylase 1 (ACC1) and SREBP1 were higher in Tg^TM4SF5^ mouse livers compared with WT mouse livers (Fig. 1E, 1F, and 1G). Molecules for hepatic lipid uptake, including *Cd36*, *Fatp2*, or *Fatp5*, were elevated in Tg^TM4SF5^ mouse livers compared with WT mouse livers, whereas molecules for fatty acid oxidation were not different (Figs. 1E and EV1A).

**Fig. 1.**
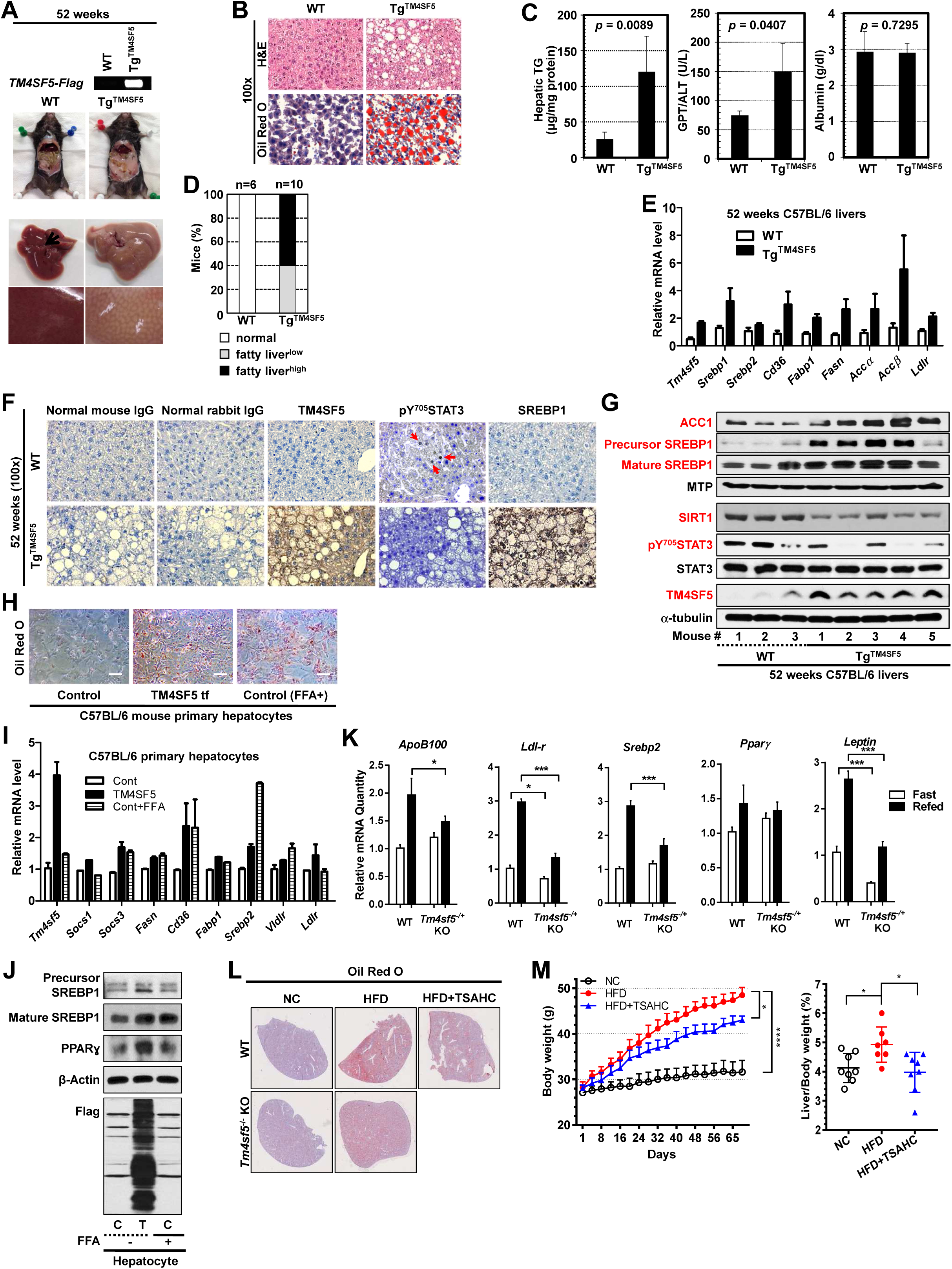
TM4SF5 overexpression caused nonalcoholic fatty liver disease. (A-B) One-year-old (52-week-old) TM4SF5-transgenic mice (C57BL/6-Tg^TM4SF5^) exhibited abdominal obesity and fatty livers (A) and lipid accumulation in livers (B) compared with wild-type (WT) mice. (C) One-year-old C57BL/6-Tg^TM4SF5^ mice exhibited enhanced hepatic triacylglyceride (TG) and alanine aminotransferase (ALT) levels but unchanged albumin levels, compared with those in WT mice. (D) Fatty liver phenotypes of C57BL/6-Tg^TM4SF5^ or WT mice. (E-G) One-year-old C57BL/6-Tg^TM4SF5^ mouse livers exhibited higher levels of mRNAs (E) and proteins (F and G) related to lipid metabolism and higher TM4SF5 and SREBP1 but lower SIRT1 and pY^705^STAT3 levels when compared with WT mouse livers. (H-I) Primary hepatocytes from C57BL/6 WT mice exhibited lipid accumulation upon TM4SF5 transfection (tf) or free fatty acid (FFA) treatment (H) and increased mRNAs related to lipid metabolism (I). (J) The transfection of TM4SF5 or treatment with FFA of primary hepatocytes resulted in increased levels of SREBP1 and PPARγ. (K) mRNAs related to lipid metabolism following refeeding in *Tm4sf5^-/+^* knockout (KO) mice that had fasted overnight, compared with those of WT mice. (L-M) Liver tissues from mice (WT and *Tm4sf5*^-/-^ KO, n≥7) on an ad lib normal chow (NC) or high-fat diet (60 kcal% fat, HFD) for 10 weeks were processed for Oil Red O staining (L) or body weight analysis (M). * *p* < 0.05, **** < 0.0001. The data represent the result of three independent experiments. Also see Figure EV1.

### TM4SF5 caused fatty acid accumulation in hepatocytes

Primary normal murine hepatocytes transfected with TM4SF5 or treated with free fatty acids (FFAs) exhibited an increased lipid accumulation (Fig. 1H) and mRNA levels of lipid metabolism-related *Srebp2* and *Cd36* (Fig. 1I). Mature SREBP1 and PPARγ protein levels also increased (Fig. 1J). Whereas refeeding of overnight-fasted WT mice increased *ApoB100*, *Ldlr*, and *Srebp2* mRNA levels in livers, but heterozygous *Tm4sf5^-/+^* mice exhibited smaller increases in the mRNAs (Fig. 1K). Lipid accumulation by a high-fat diet (HFD, 10 weeks) in WT mice was blocked by a specific TM4SF5 inhibitor TSAHC (4’-[*p*-toluenesulfonylamido]-4-hydroxychalcone(Lee et al, 2009)), whereas the livers of *Tm4sf5*^-/-^ KO mice exhibited less lipid accumulation (Fig. 1L). Body and liver/body weight gain in HFD-mice was inhibited by TSAHC treatment (Fig. 1M). A shorter period of HFD (5 weeks) was associated with TM4SF5-dependent steatosis and SREBP expression (Fig. EV1B and EV1C).

Interestingly, STAT3 phosphorylation at Tyr705 (i.e., pY^705^STAT3) was lower in Tg^TM4SF5^ mouse livers compared with WT mouse livers (Fig. 1F and 1G). Immunohistochemistry of steatotic mouse liver tissues revealed a positive correlation among TM4SF5 and SREBP1 expression and lipid droplet accumulation but a negative correlation between TM4SF5 and pY^705^STAT3 levels (Fig. 1F). TM4SF5 transfection or FFA treatment to primary hepatocytes increased the mRNA levels of *Socs3* and other lipid metabolism-related genes (Fig. 1I). STAT3 activity thus remained inactive when TM4SF5-mediated SREBP1 maturation might cause lipid synthesis and accumulation. An inverse correlation between SREBP1 maturation and STAT3 activity was downstream of TM4SF5 during NAFLD.

### TM4SF5-mediated fatty acid accumulation in adipocytes

Because adipocytes release FFAs that are taken up by hepatocytes, we also examined how TM4SF5 in 3T3-L1 adipocytes affected lipid synthesis and STAT3 activity. When TM4SF5 was suppressed in differentiated 3T3-L1 adipocytes, there were decreases in *Pparγ*, *Cd36*, *Fasn*, *Srebp1*, and *Fabp1* mRNA levels (Fig. EV2A). TM4SF5 suppression reduced lipid accumulation in 3T3-L1 cells (Fig. EV2B), suggesting that TM4SF5 expression in adipocytes also promotes lipid and FFA anabolism and accumulation. In differentiated adipocytes, STAT3 activity was negatively correlated with SREBP1 maturation (Fig. EV2C). As differentiation progressed, TM4SF5 and mature SREBP1 levels increased, whereas pY^705^STAT3 levels gradually decreased (Fig. EV2C). Thus, TM4SF5-mediated SREBP1 maturation mediates lipid synthesis and accumulation in adipocytes, which may in turn lead to lipid accumulation in hepatocytes.

### TM4SF5-mediated STAT3 activity-dependent ECM overexpression

Because one-year-old (52-week-old) Tg^TM4SF5^ mice had NAFL, we hypothesized that older mice would have more severe liver disease, resembling NASH and/or fibrosis. One-and-a-half-year-old (78-week-old) mice exhibited NASH and fibrotic phenotypes, in addition to extramedullary hematopoiesis with blood cell infiltrates in the liver (Fig. 2A and 2B). Unlike 52-week-old mouse livers, meanwhile, 78-week-old Tg^TM4SF5^ and WT mouse livers exhibited comparable levels of lipid metabolism-related genes (Figs. 1E and 2C). Molecules involved in fibrotic phenotype, immune response, and macrophage activation were dramatically elevated in 78-week-old Tg^TM4SF5^ mice compared with WT mice, indicating NASH phenotypes with inflammation (Fig. 2D). Molecular analyses of livers from older Tg^TM4SF5^ mice revealed increased pY^705^STAT3, laminin, and TM4SF5 levels, but unchanged SOCSs levels, compared with those from age-matched WT mice (Fig. 2E and 2F). The 78-week-old Tg^TM4SF5^ mice also exhibited higher collagen I, laminin, laminin γ2, and pY^705^STAT3 levels as well as active immune responses to possible pathogens and/or hepatic injury via extramedullary hematopoiesis (Fig. 2A). Notably, fibrotic livers in 78-week-old Tg^TM4SF5^ mice exhibited non-altered mature SREBP1 and elevated levels of SIRT1, compared with the fatty livers of 52-week-old Tg^TM4SF5^ mice (Fig. 2E). The TM4SF5-induced SIRT1 appeared not to positively correlate with expression levels of SREBPs and SOCSs any more in NASH-fibrotic livers. Meanwhile, immunohistochemical studies revealed the overexpression of collagen I and laminin γ2 as well as higher TM4SF5 and pY^705^STAT3 levels in 78-week-old Tg^TM4SF5^ mouse livers, compared with WT mouse livers (Fig. 2F). Interestingly, laminin γ2-positive cells were distinct from collagen I and/or α-smooth muscle actin (α-SMA)-positive cells in Tg^TM4SF5^ mouse livers (Fig. 2F).

**Fig. 2.**
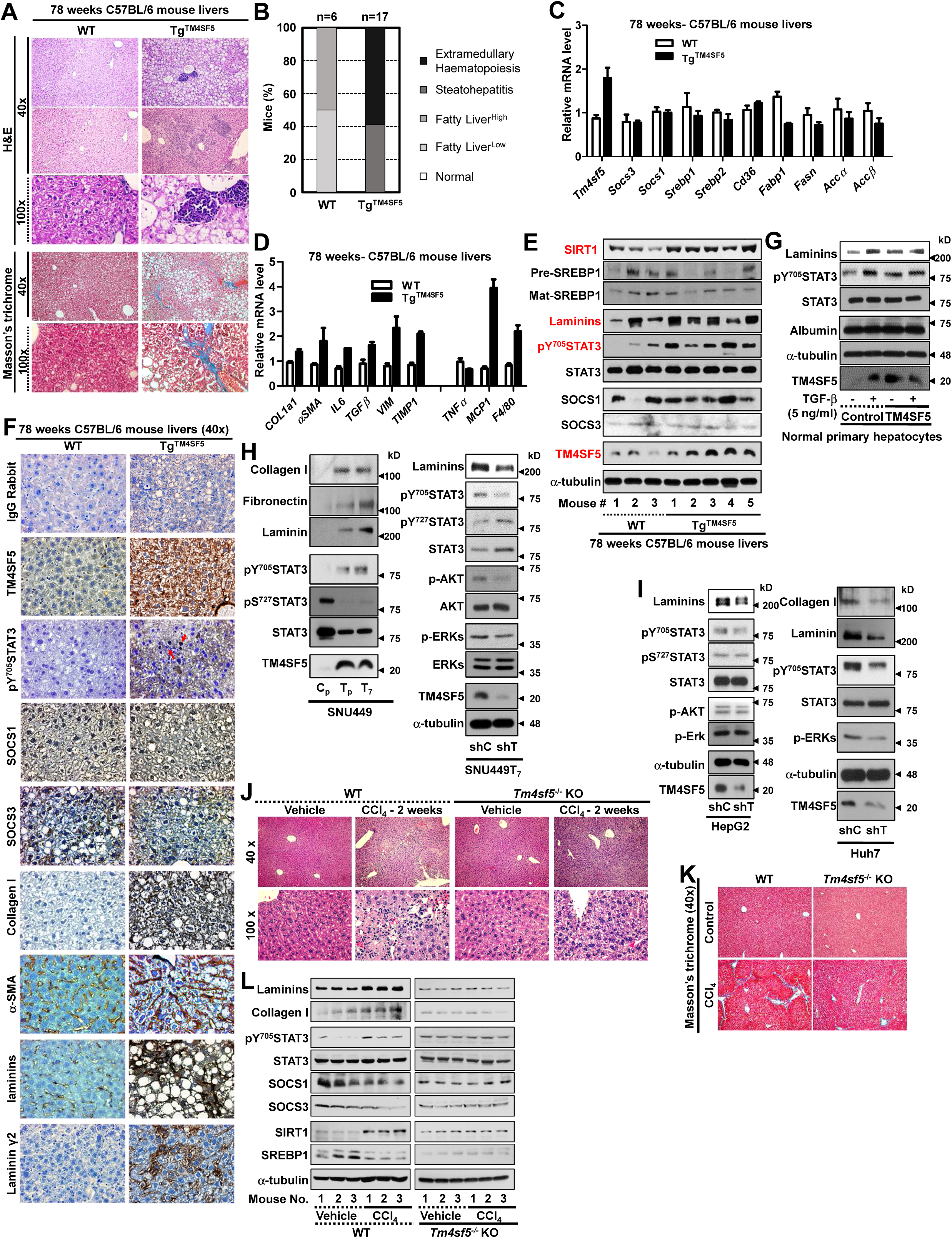
Older Tg^TM4SF5^ mice exhibited non-alcoholic steatohepatitis (NASH) and fibrotic phenotypes. (A) One-and-a-half-year-old (78-week old) C57BL/6-Tg^TM4SF5^ mice with steatohepatitic and fibrotic livers with extramedullary hematopoiesis. (B) Fatty and fibrotic liver phenotypes in 78-week old C57BL/6-Tg^TM4SF5^ and WT mice. (C-D) The livers from 78-week old C57BL/6-Tg^TM4SF5^ mice were analyzed for mRNA levels of genes related to lipid metabolism, fibrosis, immune responses, and macrophage activation. (E-F) 78-week-old C57BL/6-Tg^TM4SF5^ mouse livers were analyzed for levels of the indicated molecules and compared with WT mouse livers using western blotting (E) or immunohistochemistry (F). (G) The primary hepatocytes from WT mice were analyzed for TGFβ1-mediated laminin expression following transfection of a control or TM4SF5 plasmid. (H) Stable SNU449 cell lines ectopically expressing TM4SF5 (T_p_ and T_7_) and control cells (C_p_) were transiently transfected with shControl (shC) or shTM4SF5 (shT), prior to immunoblottings. (I) Hepatocytes endogenously expressing TM4SF5 were stably transfected with shControl (shC) or shTM4SF5 (shT), prior to immunoblotting. (J-L) The liver tissues from *Tm4sf5*^-/-^-knockout (KO) and WT mice that were administered with either vehicle or CCl_4_ for 2 weeks were analyzed via hematoxylin and eosin staining (J), Masson’s trichrome staining (K), or immunoblotting (L). The data represent the results of three isolated experiments. Also see Figure EV3.

Compared with 78-week-old WT mouse livers, 78-week-old Tg^TM4SF5^ mouse livers exhibited either non-significant changes or slightly reduced lipid metabolism-related *Srebp1/2* and *Socs1/3* mRNA levels, but elevated immune response- and macrophage activation-related collagen I (*Cola1*), *α-Sma*, *Tgfβ, Il6, Mcp1,* and *F4/80* levels (Fig. 2C and 2D). Therefore, lipid metabolism-related SREBPs and other genes and SOCSs were at most unaltered, whereas molecules for NASH or fibrosis were elevated as Tg^TM4SF5^ mice aged (>52 weeks old).

Elevated pY^705^STAT3 levels upon TM4SF5 expression (via an ectopic transfection or in the primary hepatocytes from CCl_4_-administrated mouse) were independent of further treatment with TGFβ1, IL6, or extracellular laminins, although TGFβ1 and IL6 each enhanced pY^705^STAT3 levels in cells lacking TM4SF5 expression (Fig. 2G, EV3A, and EV3B). Ligand-independent and TM4SF5-dependent STAT3 activity depended on c-SRC activity, leading to laminin expression in TM4SF5-positive hepatocytes or in the liver of CCl_4_-administrated mice (Fig. 2H, EV3A, and EV3C). The suppression of endogenous TM4SF5 in HepG2 and Huh7 cells decreased laminins and pY^705^STAT3 levels but not pS^727^STAT3 or pY^694^STAT5 levels (Fig. 2I). Following CCl_4_ treatments, *Tm4sf5*^-/-^-KO mice exhibited less macrophage infiltration and collagen I accumulation in the liver than WT mice (Fig. 2J and 2K), and did not increase SIRT1, pY^705^STAT3, or ECM levels with no changes in SOCSs levels, unlike the observations in WT mice (Fig. 2L). Thus, TM4SF5-mediated NASH/fibrosis appeared to require the TM4SF5/c-SRC/pY^705^STAT3 signaling axis.

### Alternative linkages of TM4SF5-mediated SIRT1 levels to either SREBP1 or pY^705^STAT3 during NAFLD progression

SIRTs (sirtuins) deacetylate proteins and are well-known negative regulators of genes for lipid metabolism including *SREBP1c*(Houtkooper et al, 2012). One-year-old Tg^TM4SF5^ mouse livers exhibited reduced *Sirt1* and *Sirt6* levels but enhanced *Sirt2*, *3, 4*, *5,* and *7* mRNA levels, compared with those in WT mouse livers (Fig. 3A). Fatty Tg^TM4SF5^ mouse livers showed decreased SIRT1 protein and mRNA levels, compared with WT mouse livers (Figs. 1G and 3A). TM4SF5-dependent NAFL phenotype with elevated SREBP levels may thus involve reduced SIRT1 and SIRT6 levels (Figs. 1G and 3A). TM4SF5-positive cells exhibited lower SIRT1 levels, compared with TM4SF5-negative hepatocytes, and additional palmitic acid (PA) treatment further reduced SIRT1 levels (Fig. 3B). Increased SIRT1 and decreased SREBP1 levels in the primary hepatocytes from *Tm4sf5*^-/-^ mice compared with WT mice were further decreased by PA treatment (Fig. 3B, bottom). SIRT1 transfection into the primary WT mouse hepatocytes reduced SREBP1 expression levels (Fig. 3C). *TM4SF5* and *SIRT1* gene expression tend to be increased (*p*=0.121) and decreased (*p*=0.251), respectively, whereas *SREBP1* gene expression is significantly increased (*p*=0.001) in healthy obese people compared with steatosis patients (Fig. 3D). Thus, *TM4SF5* and *SIRT1* appeared inversely related in the development of *SREBP1*-involved steatosis. Meanwhile, *Socs1* and *Socs3* mRNA levels were elevated in 52-week-old Tg^TM4SF5^ mouse livers compared with WT mouse livers (Fig. 3E), indicating SIRT1-mediated SOCS1/3 reduction (Fig. 3C). *Tm4sf5*^-/+^ mouse livers also exhibited lower SOCS1 and SOCS3 mRNA and protein levels compared with WT mouse livers (Fig. 3F and 3G), and fatty Tg^TM4SF5^ mouse livers exhibited elevated SOCS1 and SOCS3 levels (Fig. 3H), thereby leading to lower pY^705^STAT3 levels in 52-week-old Tg^TM4SF5^ mouse livers (Fig. 1G).

**Fig. 3.**
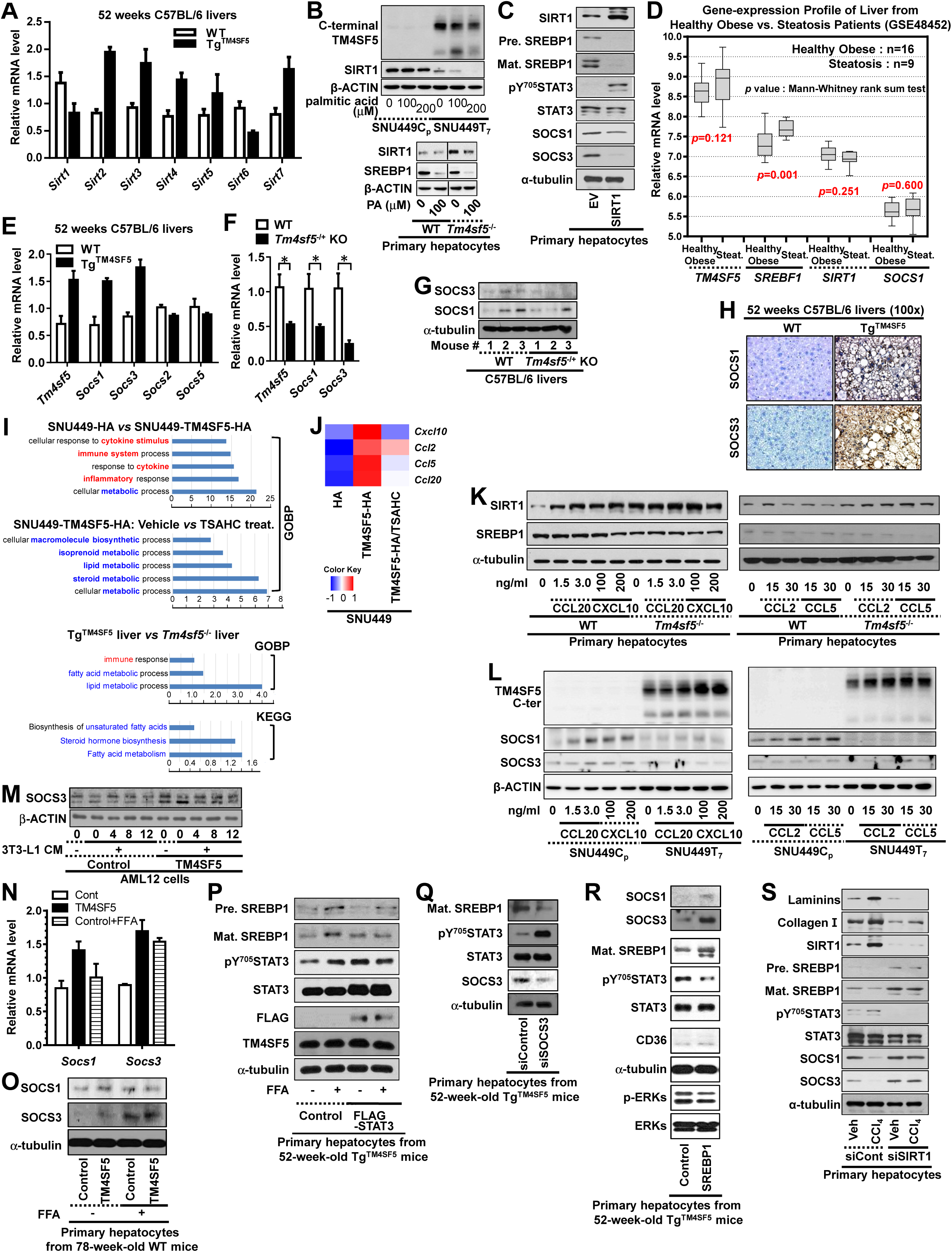
Alternative TM4SF-mediated SIRT1 linkage to either SREBP1 or STAT3 during progression to nonalcoholic fatty liver disease. (A) One-year (52-week)-old C57BL/6-Tg^TM4SF5^ mouse livers exhibited different levels of *Sirts* mRNAs compared with those of wild-type (WT) mouse livers. (B) SNU449C_p_, SNU449T_7_, or primary hepatocytes from WT or *Tm4sf5*^-/-^ mouse livers were treated without or with palmitic acid (PA) prior to immunoblotting. (C) Normal primary hepatocytes were transfected with empty vector (EV) or murine SIRT1 plasmids for 48 h, prior to immunoblotting. (D) GSE48452 gene expression profiles showed an increase in *SREBP1* (*p*=0.012) mRNA but a decrease in *SIRT1* (*p*=0.246) in the *TM4SF5-*increased (*p*=0.019) steatosis patient group, compared with a healthy obese group. (E) One-year-old C57BL/6-Tg^TM4SF5^ mouse livers exhibited higher *Socs1* and *Socs3* mRNAs levels compared with WT mouse livers. (F-G) *Tm4sf5^-/+^* heterozygous knockout (KO) mouse livers were analyzed for *Socs1* and *Socs3* mRNA (F) and protein levels (G). * *p* < 0.05. (H) One-year-old C57BL/6-Tg^TM4SF5^ and WT mouse livers were processed for immunohistochemical analysis of SOCS1 and SOCS3. (I) Gene ontology for biological process (GOBP) and Kyoto Encyclopedia of Genes and Genomes (KEGG) pathway analysis of SNU449 cell lines or mouse livers without or with TM4SF5 expression and/or TSAHC treatment. (J) A heatmap of indicated mRNA overexpression in the presence of TM4SF5 expression. (K-L) Primary hepatocytes (K) or SNU449 cell lines (L) were treated with chemokines at the indicated concentrations for 24 h, prior to immunoblotting. (M) AML12 murine hepatocytes were transfected with TM4SF5 and/or treated with conditioned media (CM) of 3T3-L1 adipocytes prior to immunoblotting. (N-O) Primary hepatocytes from normal C57BL/6 WT mice were analyzed for *Socs1* or *Socs3* mRNA (N) or protein levels (O) after TM4SF5 transfection or free fatty acid (FFA) treatment. (P) FFA treatment-induced SREBP1 maturation was examined after STAT3 transfection. (Q-R) The suppression of SOCS3 (Q) or the transfection of SREBP1 (R) in primary hepatocytes from one-year-old C57BL/6-Tg^TM4SF5^ mice was conducted prior to immunoblotting. (S) Normal primary hepatocytes were transfected with siRNA against a control sequence or *Sirt1* for 48 h, prior to immunoblotting. The data represent the results of three independent experiments.

RNA-Seq data analysis using TM4SF5-negative and -positive samples (SNU449C_p_ versus SNU449T_7_ cells and WT mouse livers versus Tg^TM4SF5^ mouse livers) resulted in changes in the expression of 16,257 (cellular samples) or 24,533 (liver tissue samples) genes after data quality check. They included genes for immune- and lipid-metabolism processes (Fig. 3I), including CCL2, CCL5, CCL20, and CXCL10. Their expressions were enhanced by TM4SF5 expression and reduced by TSAHC treatment (Fig. 3J). The treatment of CCL20 or CXCL10 to primary hepatocytes led to increased SIRT1 levels in WT primary hepatocytes, whereas *Tm4sf5*^-/-^ hepatocytes exhibited higher basal levels without any further increases (Fig. 3K). Furthermore, the treatment of the chemokines into SNU449C_p_ or SNU449T_7_ cells increased TM4SF5 and reduced SOCS1/3 levels in TM4SF5-positive SNU449T_7_ cells (Fig. 3L). Murine AML12 hepatocytes exogenously expressing TM4SF5 also had higher SOCS3 levels than control mock-transfected cells; these higher levels were maintained after treatment with conditioned media (CM) from differentiated 3T3-L1 cells (Fig. 3M), supporting the adipocyte effects on SOCS-dependent STAT3 activity in hepatocytes. TM4SF5 transfection or FFA treatment increased *Socs1* and *Socs3* mRNA and their protein levels in primary WT hepatocytes from 78-week-old mice (Fig. 3N and 3O). FFA treatment-mediated SREBP1 protein maturation in TM4SF5-expressing hepatocytes was no longer observed when the cells were transfected with STAT3 to cause pY^705^STAT3 (Fig. 3P). SOCS3 suppression in primary hepatocytes prepared from 52-week-old Tg^TM4SF5^ mice increased pY^705^STAT3 levels and reduced SREBP1 levels (Fig. 3Q), and SREBP1 overexpression increased SOCS1 and SOCS3 expression, leading to reduced pY^705^STAT3 levels (Fig. 3R). These indicate a bidirectional feedback linkage between mature SREBP1 and SOCSs under TM4SF5 expression. Indeed, patients with steatosis who exhibit higher *SREBP1* mRNA levels, compared with healthy obese individuals, also exhibited insignificantly (*p*=0.600) enhanced mean *SOCS1* mRNA levels (Fig. 3D). CCl_4_-mediated reduction in matured SREBP1 and SOC1/3 levels in the primary hepatocytes from mice treated without or with CCl_4_ for 4 weeks were abolished by additional siSIRT1 transfection, with becoming independent of CCl_4_ treatment (Fig. 3S). No CCl_4_-mediated ECM production was observed upon siSIRT1 transfection (Fig. 3S). Together these observations suggest that TM4SF5 decreased SIRT1 and increased SOCS1/3 expression to enhance SREBP levels and to inactivate STAT3, respectively, during TM4SF5-dependent NAFL, whereas TM4SF5 alternatively increases SIRT1 and restricts SOCSs levels to cause STAT3 activation, presumably leading to fibrotic ECM production.

### Clinical and animal models for TM4SF5-dependent SIRT1 for STAT3 activation and differential ECM expression

We further examined whether expression levels of TM4SF5-related molecules were relevant to liver disease status of clinical patients. Therefore, liver tissues from clinically-diagnosed nonalcoholic fatty liver (NAFL) or steatohepatitis (NASH) patients with fibrosis at different scores were immunoblotted for TM4SF5 and TM4SF5-related molecules. TM4SF5, CCL2, CCL5, SIRT1, pY^705^STAT3, and laminin γ2 in NASH tissues were obviously promoted, compared with them in NAFL tissues. Collagen I and matured SREBP1 were also generally higher in NASH tissues (Fig. 4A). This observation indicates that TM4SF5-mediated expressions of the molecules were also involved in the clinical NAFL and NASH development; especially, SIRT1 expressions were obvious in NASH but not in NAFL patients, whereas TM4SF5 expressions were higher in NASH patients than slight levels in NAFL patients. To examine the importance of STAT3 activity in ECM production, CCl_4_-administered animal models were also analyzed. Compared with control livers, CCl_4_-treated mouse livers exhibited abnormal liver cell organization, with collagen I deposition at the portal areas and slight collagen I deposition between the portal areas (i.e., moderate fibrosis) after 4 weeks-treatment and intensive deposition bridging the portal areas (i.e., incomplete cirrhosis or cirrhosis) after 16 weeks-treatment (Fig. 4B). Further, collagen I and laminins were dramatically increased simultaneously with enhanced TM4SF5 and pY^705^STAT3 levels (Fig. 4C). Collagen I expression was gradually increased toward cirrhotic livers (after 16 week CCl_4_ treatment) whereas laminins appeared highly maintained even from fibrotic livers (after 4 week CCl_4_ treatment). TM4SF5 suppression in primary hepatocytes prepared from CCl_4_-treated mouse livers reduced collagen I and laminin expression and caused STAT3 inactivation (Fig. 4D). Treatment of primary hepatocytes from CCl_4_-administered mouse livers with TSAHC(Lee et al, 2009) decreased laminins and pY^705^STAT3 levels but not pS^727^STAT3, pY^694^STAT5, and albumin levels (Fig. 4E). STAT3 suppression inhibited TM4SF5-mediated ECM expression (Fig. 4F), and pharmacological c-SRC inhibition abrogated pY^705^STAT3 and ECM levels (Fig. EV3C). Whereas fibronectin (*FN1*) mRNA levels were the same in both untreated control and CCl_4_-treated mouse livers, mRNA levels of elastin and the α2, α3, α5, γ2, and γ3 laminin chains were markedly elevated in CCl_4_-treated livers (Fig. 4G). Immunohistochemistry revealed increased TM4SF5, pY^705^STAT3, α-SMA, collagen I and IV, laminin, and laminin γ2 expression levels in CCl_4_-treated livers compared with control livers (Fig. 4H). Notably, laminin and laminin γ2 expression at locally patched patterns were different from those of collagen and α-SMA, which were mostly localized at the septa or bridging areas (Fig. 4H), suggesting differential cell-types specific for the ECM expressions. Thus, TM4SF5-dependent ECM induction positively correlated with STAT3 phosphorylation during the development of fibrosis/cirrhosis phenotypes.

**Fig. 4.**
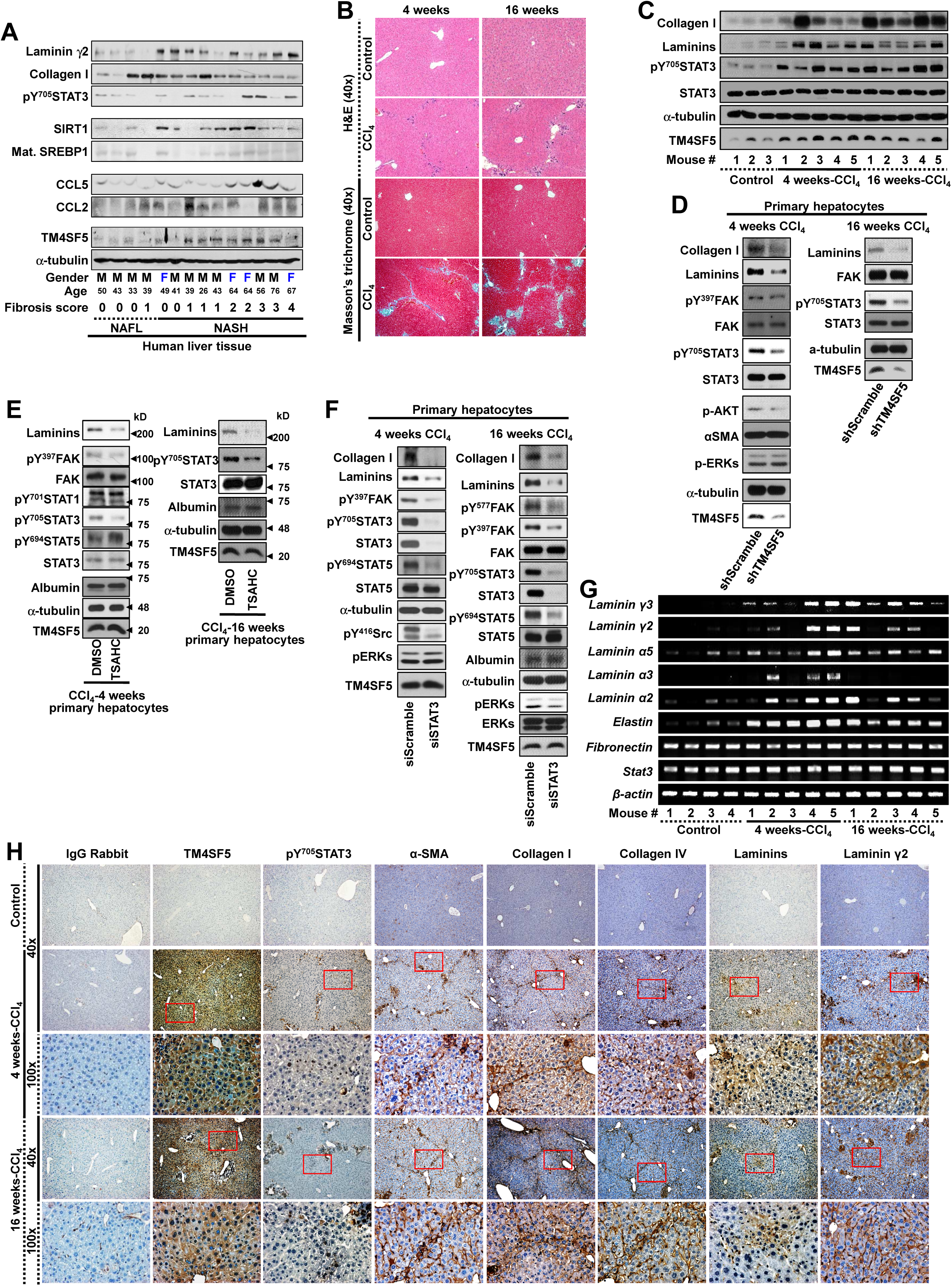
CCl_4_ administration led to TM4SF5-dependent STAT3 activation for differential ECM expression. (A) Liver tissue extracts from clinically-diagnosed NAFL and NASH patients with the indicated fibrosis scores were prepared for the immunoblotting against theindicated molecules. (B-C) Wild-type (WT) mice were treated without (control) or with CCl_4_ for 4 or 16 weeks, prior to liver analysis via hematoxylin and eosin and Masson’s trichrome staining (B) or immunoblotting (C). (D) TM4SF5 suppression in primary hepatocytes from CCl_4_-treated mice was conducted prior to immunoblotting. (E and F) TSAHC (a specific anti-TM4SF5 small compound) treatment to (E) or suppression of STAT3 (F) in primary hepatocytes from CCl_4_-treated mice was conducted prior to immunoblotting. (G-H) WT mice were treated with CCl_4_ or without (control) for 4 or 16 weeks, prior to liver analysis via reverse transcriptase polymerase chain reaction (RT-PCR) (G) or immunohistochemistry (H). The data represent the results of three isolated experiments.

### Differential TM4SF5/STAT3-dependent expression of laminins and collagen I

We next determined how hepatic ECM induction was regulated by TM4SF5. To explore which cell types might be responsible for collagen I or laminin expression, we examined the activity of the laminin γ2 (*Lamc2*) and collagen I α1 (*Col1a1*) chain promoters in AML12 murine hepatocytes and LX2 human HSCs. Collagen I α1 and laminin γ2 promoter regions were constructed with STAT3-binding elements upstream of the luciferase gene sequence (Fig. 5A). The effect of TM4SF5 or STAT3 expression on the transcriptional activation of the *Lamc2* promoter was greater in AML12 hepatocytes than in LX2 HSCs, whereas the effect on the *COL1A1* promoter was much greater in LX2 HSCs than in AML2 hepatocytes (Fig. 5B). *Lamc2* appeared to be preferentially expressed in hepatocytes, whereas *Col1a1* was preferentially expressed in HSCs. Laminin immunostaining in mouse liver tissue overlapped with certain cells that were also positive for TM4SF5 (Fig. 5C). The TM4SF5/STAT3 signaling axis in hepatocytes might thus have a greater functional impact on *Lamc2* transcription.

**Fig. 5.**
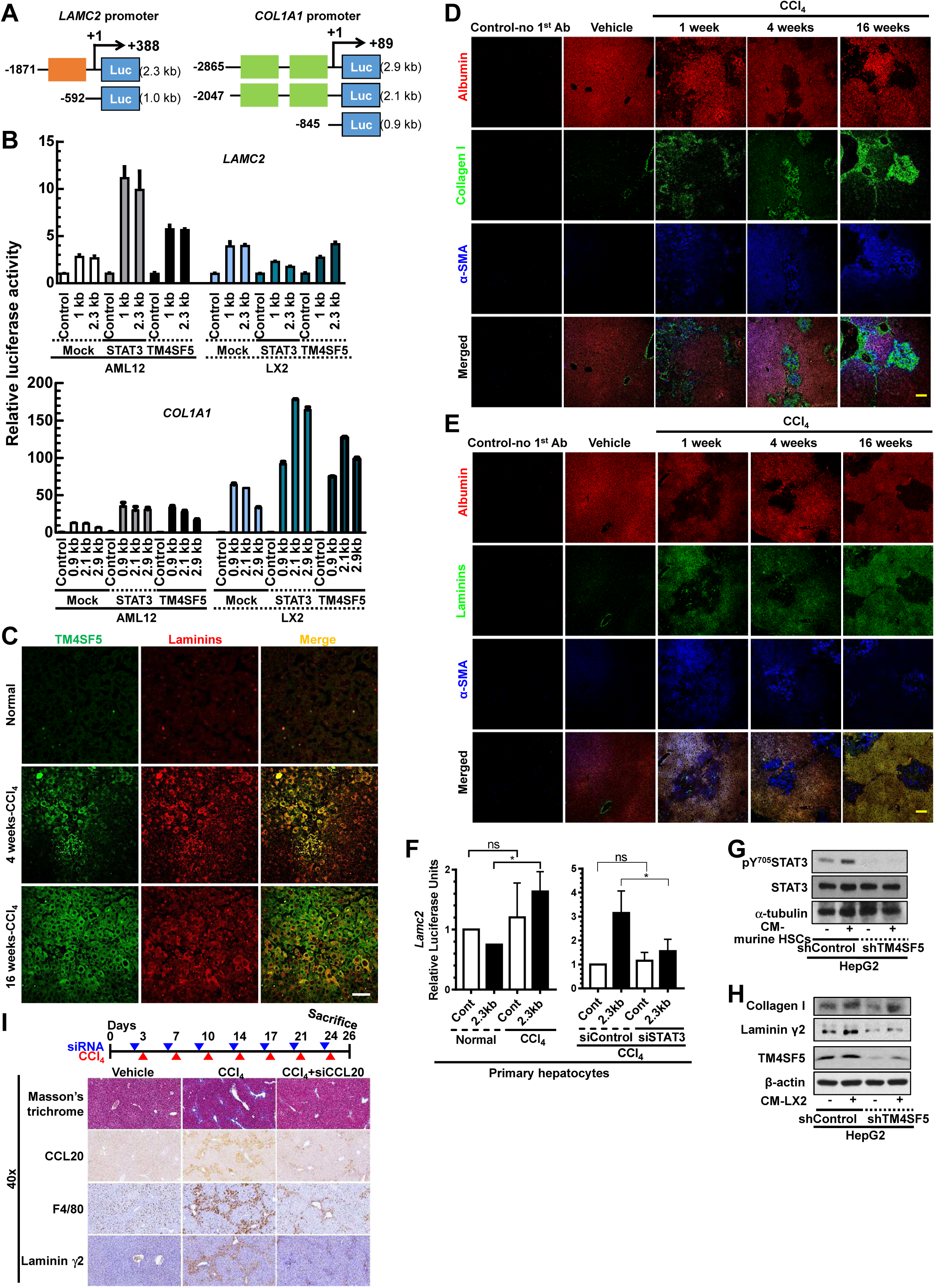
TM4SF5/STAT3-dependent laminin and collagen I expression in hepatocytes and hepatic stellate cells, respectively. (A) Luciferase constructs with STAT3-responsive sequences upstream of the laminin γ2 (*LAMC2*) or collagen I α1 (*COL1A1*) gene. (B) The transfection of TM4SF5 or STAT3 into AML12 murine hepatocytes or LX2 human hepatic stellate cells (HSCs) was conducted prior to the extracellular matrix (ECM) promoter (*LAMC2* or *COL1A1* promoter) activity analysis. (C) Laminin and TM4SF5 were co-stained in CCl_4_-treated mouse livers. (D-E) Immunostaining of collagen I and laminin in CCl_4_-treated mouse livers was conducted in parallel to α-SMA and albumin staining. Scale bars, 100 μm. (F-G) HepG2 cells with or without TM4SF5 suppression were treated without (-) or with (+) the conditioned media (CM) of primary murine HSCs (F) or LX2 (G), prior to immunoblotting. (H) Masson’s trichrome staining or immunostaining was conducted on control- or CCl_4_-treated mouse livers without or with intraperitoneal (IP) injections of si*Ccl20*. Magnification: 40×. The data represent the results of three independent experiments.

When mice were treated with CCl_4_, collagen I expression overlapped with α-SMA expression indicating activated HSCs, whose pattern was negatively superimposed with albumin staining to indicate hepatocytes (Fig. 5D). Meanwhile, laminin expression positively overlapped with albumin immunostaining and negatively overlapped with α-SMA staining after CCl_4_ treatment for 4 or 16 weeks. After a short (1-week) CCl_4_ treatment, both α-SMA and albumin immunostaining overlapped with laminin staining, but the overlapping albumin and laminin staining became more intense at later time points (Fig. 5E). Primary hepatocytes from normal or CCl_4_-treated mice showed a significant activation of *Lmac2* promoter, which was abolished by STAT3 suppression (Fig. 5F), indicating STAT3 activity in hepatocytes could be important for laminin γ2 expression during CCl_4_-induced fibrosis. CM collected from primary murine HSC culture was used to treat HepG2 cells expressing endogenous TM4SF5 that had been pre-transfected with shRNA against a control sequence or TM4SF5. pY^705^STAT3 in HepG2 cells was enhanced by mouse HSC-CM treatment, which was blocked by TM4SF5 suppression (Fig. 5G). TM4SF5 suppression in HepG2 cells also abolished laminin γ2 expression depending on the human HCS (LX2)-CM treatment. Meanwhile, collagen I expression in HepG2 cells was also promoted by the HSC-CM treatment, and TM4SF5 suppression caused collagen I expression to be more dependent on the CM treatment (Fig. 5H). Moreover, the suppression of CCL20 decreased macrophage infiltration, leading to reduced collagen I and laminin γ2 expression for less fibrotic phenotypes (Fig. 5I). The characteristics of laminin expression thus differed from those of collagen I during the development of fibrosis/cirrhosis phenotypes.

### Laminin γ2 suppression blocked TM4SF5-mediated liver injury and fibrosis

We next examined the laminin expression in clinical human liver tissues. Immunostaining of control, peritumoral, and tumor liver tissue from HCC patients revealed elevated TM4SF5, laminins, collagen I, and pY^705^STAT3 levels in peritumoral (presumably with NASH and fibrosis phenotypes) and tumor regions, compared with control regions (Fig. 6A). Laminin-positive cells again appeared to be hepatocytes with large cytosols, and were different from collagen I-positive cells (Fig. 6A). Analysis of publically available data also showed a positive correlation between enhanced *TM4SF5* and LAMININ γ2 (*LAMC2*) expression in cirrhosis, compared with healthy human liver tissues (Fig. EV4). *TM4SF5* (*p*<0.001) mRNA expression as well as *COL1A1* (*p*<0.001) and *LAMC2* (*p*=0.015) mRNA expression were also higher in patients with cirrhosis compared with healthy individuals, whereas *SREBP1* (*p*> 0.222) mRNA levels were not differential, and *SOCS1* (*p*=0.065) mRNA levels were lower (Fig. EV4A and EV4B). Thus, the development of pre-cancerous cirrhosis appeared to involve the expression of *LAMC2* as well as *COL1A1*.

**Fig. 6.**
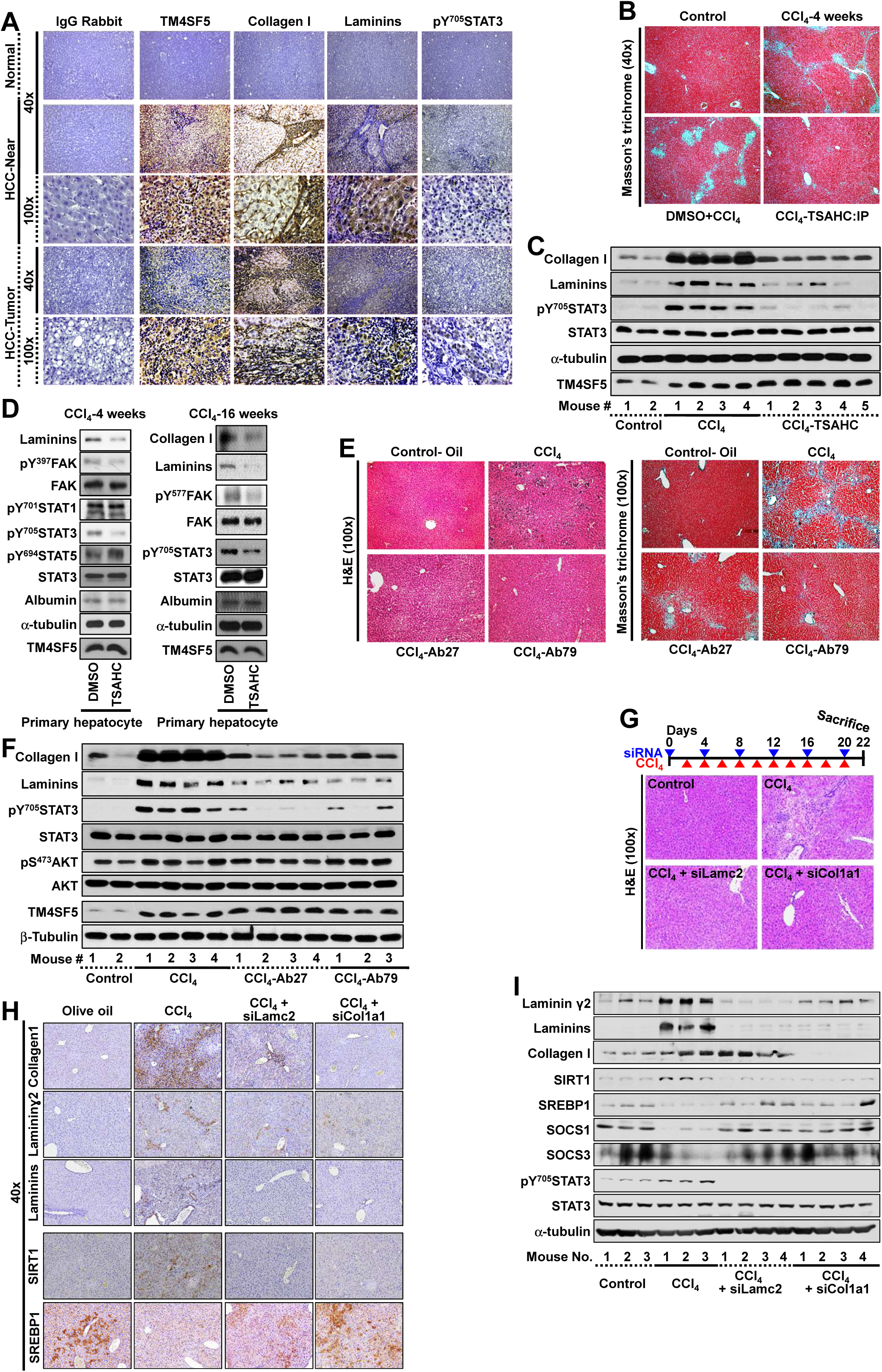
Laminin γ2 suppression blocked TM4SF5-mediated liver injury and fibrosis. (A) Clinical human liver cancer tissues, cancer-adjacent tissues presumably containing fibrotic/cirrhotic lesions, and healthy liver tissues were analyzed for levels of the indicated molecules. (B-D) Mice were treated with CCl_4_ for 4 weeks in the absence or presence of intraperitoneal (IP) treatment of DMSO or TSAHC. Liver tissues were stained for collagen I (B) and processed for immunoblotting (C). (D) Primary hepatocytes from the livers of mice treated with CCl_4_ for 4 or 16 weeks were treated in the presence of DMSO or TSAHC for 24 h, prior to immunoblotting. (E-F) Mice were treated with CCl_4_ for 4 weeks in the absence or presence of intraperitoneal (IP) injection of anti-TM4SF5 antibody (Ab27 and Ab79). Liver tissues were processed for hematoxylin and eosin or Masson’s trichrome staining (E) or immunoblotting (F). (G-I) The suppression of either laminin γ2 or collagen I α1 was conducted in BALB/c mice without or with CCl_4_ as indicated. Liver tissues were processed for hematoxylin and eosin staining (G), immunostaining (H), and immunoblotting (I). The data shown represent the results of three independent experiments. Also see Figure EV4.

Collagen I deposition in CCl_4_-treated fibrotic livers was blocked after intraperitoneal treatment with TSAHC but not with control vehicle, leading to a more normal-like liver phenotype (Fig. 6B). The TSAHC-mediated inhibitory effects on fibrosis/cirrhosis were accompanied by decreases in CCl_4_-induced collagen I and laminin expression and pY^705^STAT3 levels (Fig. 6C). Treatment with TSAHC of primary mouse hepatocytes prepared from CCl_4_-treated mouse livers decreased ECM and pY^705^STAT3 levels, and focal adhesion kinase (FAK, a TM4SF5-downstream effector(Jung et al, 2012)) activity (Fig. 6D). These CCl_4_-induced ECM and pY^705^STAT3 levels also were blocked by treatment with chimeric anti-TM4SF5 monoclonal antibodies (i.e., Ab27 and Ab97), although TM4SF5 levels were unchanged (Fig. 6E and 6F). Thus, anti-TM4SF5 reagents could inhibit STAT3 activity and ECM production (collagen I as well as laminin γ2) during CCl_4_-induced fibrosis/cirrhosis.

The significance of TM4SF5-mediated laminin γ2 expression in fibrosis development was examined by tail vein injection of siLamc2 during CCl_4_ administration. The pre-administration of siRNA against laminin γ2 (*Lmac2*) or the collagen α1 (*Col1a1*) chain before CCl_4_ treatment (for 4 days) resulted in less liver damages, compared with the disordered liver phenotypes caused by CCl_4_ treatment alone (Fig. EV5A). These reduced levels of liver damage correlated with decreased laminin and pY^705^STAT3 levels, even when the expression of either laminin γ2 or the collagen α1 chain was suppressed (Fig. EV5B). Laminin expression was reduced by the suppression of either laminin γ2 or collagen α1 chain (Fig. EV5B), presumably indicating a global effect among ECM proteins. *Lamc2* suppression for a longer period blocked CCl_4_-mediated immune-cell infiltrates and hepatic damages (Fig. 6G), ECM deposition and SIRT1 expression (Fig. 6H), and pY^705^STAT3 and ECM expression levels (Fig. 6I), as did *Col1a1* suppression. Interestingly, CCl_4_-mediated changes in SIRT1, SREBP1, and SOCS expressions were reversed by either *Lamc2* or *Col1a1* suppression (Fig. 6I). Taken together, the expression profile of laminins could differ from that of collagen I during, and laminins and collagen I were primarily expressed in hepatocytes and HSCs, respectively, during NASH with fibrosis/cirrhosis.

## Discussion

This study shows that TM4SF5-mediated modulation of SIRT1 expression can promote chronic exacerbation of liver pathologies. TM4SF5 down-regulated SIRT1 expression, leading to upregulated SREBPs for fatty acid synthesis and accumulation during NAFL, whereas SOCSs levels concomitantly remained high for STAT3 inactivation. As the severity of disease progressed in the presence of the TM4SF5-mediated inflammatory environment, TM4SF5 upregulated SIRT1 to restrict SREBPs and SOCSs expression leading to ligand-independent STAT3 activation for ECM production during NASH and fibrosis. These observations were valid in *TM4SF5*-transgenic, diet- or chemical-induced animal models; clinical liver tissues from human patients; and public GSE datasets. Furthermore, TM4SF5 expression increased collagen I expression in HSCs and laminin/laminin γ2 in hepatocytes, respectively, upon STAT3 activation. Anti-TM4SF5 antibody, TSAHC treatment, and laminin γ2 or CCL20 suppression all abrogated CCl_4_-treated liver damage and fibrotic phenotypes. TM4SF5 expression in hepatocytes may thus play dynamic roles in signal transduction leading to chronic progressions of such liver diseases (Fig. 7).

**Fig. 7.**
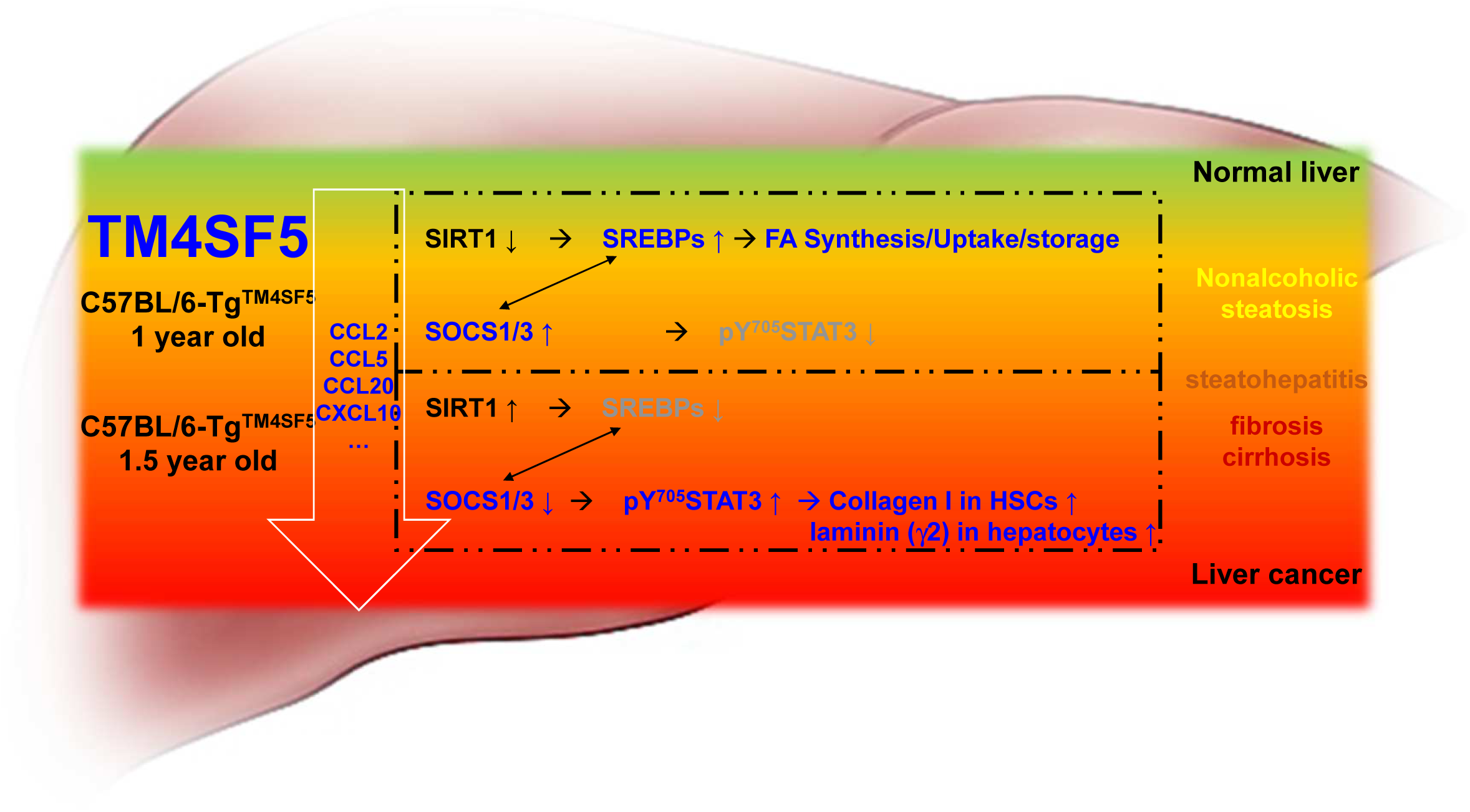
Schematic modeling of the TM4SF5-mediated progression of liver disease. TM4SF5-mediated signaling pathways in primary cell and animal models resulted in the development of chronic and multiphasic liver disease, including steatosis, steatohepatitis, and fibrosis/cirrhosis. During nonalcoholic steatosis, TM4SF5 decreased SIRT1 and thereby increased SREBP1, leading to fatty acid accumulation in livers, higher SOCS levels, and lower STAT3 activity. In older animals, TM4SF5 led to increased SIRT1 via actions by TM4SF5-dependent pro-fibrotic cytokines and chemokines, leading to reduced SREBP1 and SOCSs levels, which in turn resulted in STAT3 activation and differential ECM expression in livers (i.e., collagen I expression in HSCs and laminin expression in hepatocytes). Pre-cancerous phenotypes in livers appeared to be associated with a positive correlation between SREBP1 and SOCS in a bidirectional manner; the upregulation or downregulation of these proteins appeared to depend on TM4SF5 expression and the inflammatory environment.

The direction of modulation of SIRT1 expression levels by TM4SF5 depended on the pathological status of the liver. The TM4SF5-mediated inflammatory environment with pro-fibrotic cytokines and chemokines may alternatively transduce signaling to and/or influence on either the SIRT1^reduced^/SREBPs^increased^ or SIRT1^increased^/SOCSs^reduced^/pY^705^STAT3^increased^ axis. TM4SF5 promoted SREBPs expression in fatty livers via the down-regulation of SIRT1, which deacetylates and negatively regulates SREBPs activity and expression(Agarwal & Agarwal, 2017). SREBP-1a and SREBP-1c or SREBP-2 are involved in FFA or cholesterol synthesis, respectively(Moon, 2017). SIRT1 overexpression decreased mature SREBP1 and SOCS1/3 levels. Here, SOCS suppression decreased mature SREBP1, and SREBP1 overexpression promoted SOCS1/3 levels. These findings thus support a (bidirectional) positive crosslink between SREBP1 and SOCS1/3 (Fig. 3). Together with the inverse changes in SIRT1 and SREBPs downstream of TM4SF5, the levels of pY^705^STAT3 (not pS^727^STAT3) or SOCS1/3 were thus negatively or positively correlated with SREBP levels, respectively. When liver tissues of NAFL and NASH (with fibrosis) patients were compared, SIRT1 expression was dramatically increased in NASH patients, whereas TM4SF5 expressions were slight and further enhanced in NAFL and NASH patients, respectively. Thus, TM4SF5-mediated SIRT1 expression might depend on physiological factor(s) for NASH, that is inflammation. The inflammatory environment during NASH/fibrosis following steatosis may alternatively promote SIRT1 expression and reduced SOCS for STAT3 activation in TM4SF5-dependent manners. Chemokines and cytokines may serve as a link between inflammation, pre-cancerous fibrosis/cirrhosis, and liver cancer(Biancheri et al, 2014; Marra & Tacke, 2014). IL6, IL8, TGFβ1, CCL2, CCL5, CCL20, or CXCL10 are involved in NAFLD/NASH and fibrosis(Marra & Tacke, 2014). More specifically, proinflammatory CCL5 expression in hepatocytes during steatosis contributes to the development of early-stage liver fibrosis without significant impacts on steatosis(Li et al, 2017). Additionally, we found that CCL2, CCL5, CCL20, and CXCL10 expressions were increased in TM4SF5-positive cells and liver tissues. CCL2, CCL5, CCL20, or CXCL10 further increased TM4SF5 expression and decreased SOCS1/3 levels in TM4SF5-positive cells (but not in TM4SF5-negative cells), presumably as their pro-fibrotic roles upon TM4SF5 expression. Thus, TM4SF5-dependent inflammatory cytokines/chemokines decreased SOCS3 levels for active STAT3-mediated ECM production and also induced TM4SF5 to activate STAT3 downstream of active c-Src(Jung et al, 2005) that directly binds TM4SF5(Jung et al, 2013).

Here, TM4SF5- and STAT3-dependent laminin expression in hepatocytes was important for fibrotic phenotypes; laminin γ2 or collagen I suppression was sufficient to block liver injury and fibrotic phenotypes. Although *in vitro* hepatocytes undergo an epithelial-mesenchymal transition in the presence of TGFβ1, leading to fibrotic phenotypes(Zeisberg et al, 2007), *in vivo* analyses have not revealed a role of hepatocytes in liver fibrosis(Taura et al, 2010). HSC-produced collagen I plays an important role in liver fibrosis(Yang et al, 2014). However, in this study, active STAT3 induced laminin and laminin γ2 expression, presumably in hepatocytes, whereas collagen I in HSCs. A certain proportion of patients with NASH and fibrosis (from 2.4% to 12.8% with or without cirrhosis) will sequentially develop HCC(Anstee et al, 2019). Thus, laminin/laminin γ2 expression during fibrosis/cirrhosis may provide an environment in which hepatocytes can proliferate uncontrollably during hepatic carcinogenesis. There is also considerable immunostaining for laminin γ2 in the cytoplasm of hepatic carcinoma cells at the invasive front of a tumor(Aishima et al, 2004), and laminin γ2 is a novel serum biomarker for HCC(Kiyokawa et al, 2017). Thus, the modulation of laminin γ2 expression in hepatocytes may be a promising target for preventing pre-cancerous liver disease and HCC. It is thus interesting that anti-TM4SF5 reagents (TSAHC and anti-TM4SF5 antibody) as well as siRNAs against *Ccl20* or *Lamc2* could block TM4SF5-mediated liver disease.

Together, our findings indicate that TM4SF5 plays important roles in the progression of multiphasic liver disease via the TM4SF5-mediated SIRT1/SREBPs or SIRT1/SOCSs/STAT3 signaling axis and via TM4SF5/STAT3-mediated ECM induction in hepatocytes or HSCs. Thus, TM4SF5 and downstream effectors, including CCL20, SIRT1 and laminin γ2, are promising therapeutic targets for the treatment of liver disease.

## Materials and Methods

### Cells

TM4SF5-lacking control (hepatocarcinoma SNU449 or SNU761 cells and normal murine hepatocyte AML12) or TM4SF5-expressing (endogenously expressing HepG2 and Huh7 or ectopically expressing SNU449Tp or SNU449T_7_) cells have been described previously (Lee et al, 2008). Human hepatic stellate cells (HSCs, LX2) were cultured in DMEM-H (WelGene, Daegu, Republic of Korea) as previously described(Kang et al, 2012a). Cells were purchased from either the Korean Cell Bank (Seoul, Korea) or ATCC (Manassas, VA, USA). Stable cells were maintained in RPMI-1640 (WelGene) containing 10% fetal bovine serum (FBS), G418 (250 μg/ml), and antibiotics (Invitrogen, Grand Island, NY, USA). Cells were passaged every 3-4 days at a ratio provided by the manufacturer. Cells were monitored for mycoplasma contamination.

### Transfection

siRNA, shRNAs, or cDNA plasmids were transiently transfected for 48 h using Lipofectamine RNAiMAX or Lipofectamine 3000 (Thermo Fisher Scientific, Waltham, MA, USA), according to the manufacturer’s protocols.

### Mice

TM4SF5 transgenic mice were generated (Macrogen, Seoul, Korea) and the expression of TM4SF5 was confirmed(Kang et al, 2012a). pcDNA3-TM4SF5-FLAG was digested with *NruI* and *DraIII*. The fragments consisting of the CMV promoter, whole TM4SF5-FLAG sequence, and BGH polyA region were purified and microinjected into fertilized eggs from C57BL/6 mice according to standard procedures. Two-week-old founders were screened by PCR using the primer set for CMV-F1 (forward primer, 5’-CGCTATTACCATGGTGATGCG-3’) and TM4SF5-R1 (reverse primer, 5’-AGACACCGAGAGGCAGTAGAT-3’). TM4SF5-knockout (*Tm4sf5*^-/-^) mice were generated via embryo injection and transferred to normal healthy female C57BL/6 mice (Macrogen) by targeting the exon 1 or exons 3-5 regions coding the second (long) extracellular loop (LEL or EC2) in the mouse *Tm4sf5* gene, using *Tm4sf5* genes digested by Cas9 protein and sgRNAs (i.e., RG1 and RG2 for exon 3 up and down, respectively; RG3 for the intron between exon 4 and 5; and RG4 for exon 5). The identification of the deletion mutation in F0 founders was performed using a T7E1 assay on tail genomic DNA to identify the heteroduplex formation between WT and mutant PCR products (Macrogen). Genotyping was performed using primers for mouse-*Tm4sf5* (forward 5′-GTAGTATGCGGGAGGCACTG-3′, reverse 5′-GGGTGACCACTCAGACTTCC-3′). *Tm4sf5^-/+^* heterozygotes from the F1 litter mates were bred to generate *Tm4sf5^-/-^* homozygotes.

### CCl_4_-treated animal models

Four-week-old mice (BALB/c) were purchased from Orient. Co. Ltd (Seungnam, Korea). C57BL/6 wild-type or *Tm4sf5^-/-^* KO mice were used. The mice were housed in a specific pathogen-free room with controlled temperature and humidity. All animal procedures were performed in accordance with the procedures of the Seoul National University Laboratory Animal Maintenance Manual and with IRB approvals from the Institute of Laboratory Animal Resources Seoul National University. Five-week-old mice (n ≥ 5) were injected intraperitoneally with or without CCl_4_ (Sigma-Aldrich, St. Louis, MO, USA; 1 mg/kg, 3 times/week) in 40% olive oil. 4’-(*p*-toluenesulfonylamido)-4-hydroxychalcone (TSAHC) (Lee et al, 2009) (50 mg/kg in 40% DMSO) or of the Ab27 (5 µg/mouse) or Ab79 (16 µg/mouse) anti-TM4SF5 antibodies were injected intraperitoneally for 4 weeks on the day after each CCl_4_ administration. After 1, 2, 4, or 16 weeks, the mice were euthanized with ether, the tissues were resected, and one piece of tissue was immediately frozen in liquid N_2_ while a second piece was embedded in paraffin, or alternatively used for primary hepatocyte preparation. Mouse tail veins were injected with siRNA (Amibon: Silencer™ Pre-Designed siRNA, In-Vivo Ready) against mouse laminin γ2 (*Lamc2*, Assay ID 100303), collagen I α1 (*Col1a1,* Assay ID 160310) at 3 mg/kg, or CCL20 (*Ccl20*, Assay ID 72932) at 1.66 mg/kg in phosphate buffered saline (PBS) one day before each intraperitoneal CCl_4_ administration (5 mg/kg/injection) for the indicated schedules.

### High-fat-diet animal models

Four-week-old wild-type or knock-out C57BL/6 mice (n ≥ 7) were maintained on an ad lib normal chow or high-fat diet (60 kcal% fat, Orient. Co. Ltd) for 5 or 10 weeks without or with intraperitoneal treatments of TSAHC (Lee et al, 2009) at a 2 ∼ 3 day interval. Body weights were frequently recorded.

### Primary cell preparation

Primary hepatocytes were isolated from 4-6-week-old BALB/C mice (Orient. Co. Ltd.) or 52- or 78-week-old C57BL/6 mice or C57BL/6-Tg^TM4SF5^ mice by perfusion of the liver using collagenase type II (Life Technologies, Carlsbad, CA, USA) (Ateş et al, 2012). The hepatocytes were cultured in William’s E Medium (Life Technologies) supplemented with 10% FBS (GenDEPOT, Barker, TX, USA) and primary hepatocyte maintenance supplements (Life Technologies) on collagen-precoated plates. For the isolation of hepatic stellate cells (HSCs), non-parenchymal sufficient supernatant was centrifuged using 50%/30% Percoll (Sigma-Aldrich). The top layer containing HSCs was cultured in RPMI 1640 (WelGene) medium with 10% FBS on collagen-precoated plates.

### Co-culture

Cells were co-cultured using cell culture inserts (SPLInsert^™^, SPL Life Sciences, Pocheon-si, Korea) with 0.4 μm pores to separate the cell populations. Primary hepatocytes transiently transfected for 24 h with control scrambled shRNA or shTM4SF5 were plated on the bottom chamber under serum-free condition and primary HSCs were plated in the insert. After co-culturing for 24 h, the cells were harvested for western blotting.

### Western blotting

Subconfluent cells in normal culture media, cells transiently transfected for 48 h with control or specific siRNA or shRNA against the indicated molecules, or liver tissues from human patients or animal models were harvested for whole cell or tissue extracts using a modified RIPA buffer as described previously (Lee et al, 2008; Ryu et al, 2014). Cells were treated with human or murine cytokines or chemokines (BioLegend, San Diego, CA) for 24 h prior to cell extract preparation. The primary antibodies included those against pY^397^FAK, SOCS1, CD36, or laminins (Abcam, Cambridge, UK); pY^416^SRC, AKT, p-ERKs, ERKs, pY^694^STAT5, pS^727^STST3, FLAG, or pY^701^STAT1 (Cell Signaling Technology, Danvers, MA, USA); pY^705^STAT3 (Millipore, Billerica, MA, USA); α-tubulin, or α-SMA (Sigma-Aldrich); c-SRC, pY^577^FAK, pS^473^AKT, β-actin, β-tubulin, SOCS3, SIRT1, precursor SREBP1, matured SREBP1, PPARγ, PPARα, MTP (microsomal triglyceride transfer protein), STAT5, or laminin γ2 (Santa Cruz Biotechnology, Santa Cruz, CA, USA); FAK (BD Transduction Laboratories, Bedford, MA, USA); STAT3 (Millipore, Solna, Sweden); collagen I (Acris Antibodies, Herford, Germany); or TM4SF5 (Lee et al, 2008).

### Study approval

Human liver tissues were obtained after obtaining informed consent from patients with HCC who underwent surgery at the Liver Research Institute, Seoul National University Hospital (Seoul, Korea) or Ewha Womans University Mokdomg Hospital (Seoul, Korea). HCC regions (HCC-T) and pre-HCC regions (HCC-N) adjacent to the HCC regions or NAFL and NASH regions were obtained in accordance with IRB-approved protocols.

### Immunohistochemistry and tissue staining

Immunohistochemistry of mouse or human liver tissues was performed using primary antibodies for TM4SF5(Lee et al, 2008), collagen I, collagen IV (Acris Antibodies), pY^705^STAT3 (Cell Signaling Technology), normal rabbit or mouse IgG, SOCS3 (Santa Cruz Biotechnology), α-SMA (Sigma-Aldrich), SOCS1, laminins, albumin (Abcam), SREBP1, or laminin γ2 (Santa Cruz Biotechnology). The mouse liver tissues were prepared and processed for Masson’s trichrome staining, hematoxylin and eosin staining, or immunostaining as previously described(Zhong et al, 2009). Ten random images per slide were saved using a digital slide scanner (MoticEasyScan, Motic, British Columbia, Canada).

### Immunofluorescence

Liver tissues from mice treated without or with CCl_4_ were immunostained using antibodies against TM4SF5, laminins, albumin, collagen I, or α-SMA, as previously described(Ryu et al, 2014). Immunofluorescent images were acquired on a microscope (BX51TR, Olympus, Tokyo, Japan). Ten random image fields for each experimental condition were saved.

### PCR

Total RNAs from animal liver tissues or cells were isolated using TRIzol Reagent (Invitrogen), and their cDNAs were synthesized using amfiRivert Platinum cDNA synthesis master mix (GenDEPOT) according to the manufacturer’s instructions. The cDNAs were subjected to RT-PCR using the Dream Taq Green PCR master mix (Thermo Scientific, San Jose, CA, USA). Quantitative real time PCR (q-PCR) was prepared with LaboPass^TM^ EvaGreen Q Master (Cosmo Genetech, Seoul, Korea) and performed with the CFX Connect^™^ Real-Time PCR (Bio-Rad, Hercules, CA, USA). The mRNA levels were normalized against 18S rRNA, using the ddCq method. CFX Maestro^™^ software (Sunnyvale, CA, USA) was used to analyze the data. Primers were purchased from Cosmo Genetech (Seoul, Korea). The primer sequences are shown in Table S1.

### Adipocyte differentiation

Mouse 3T3-L1 preadipocytes (a kind gift from Dr. Jae Bum Kim, Department of Life Science, Seoul National University) were seeded into 6-well culture plates using DMEM supplemented with 10% newborn bovine serum (Thermo Fisher), and 1% penicillin/streptomycin at a density of 1 × 10^5^ cells/well. Two day after confluence (day 0), pre-adipocytes were treated with adipocyte differentiation medium containing 1 μM dexamethasone, 0.5 mM IBMX, and 10 μg/ml insulin (Sigma-Aldrich) for 2 days. The media then were replaced with DMEM supplemented with 10% FBS and insulin (10 μg/ml) for an additional 2 days. The differentiation level was determined on day 12.

### ECM-luciferase assay

To analyze the activity of the promoters, *Lamc2* promoters (encoding the regions from −1871 to +388 and −592 to +388) and *Col1a1* promoters (encoding the regions from −2865 to +89, −2047 to +89, and −845 to +89) were amplified by PCR and cloned into the pGL3-basic vector. LX2 or AML12 cells were seeded in 48 well plates using Lipofectamine 3000 transfection reagent (Invitrogen). One day after transfection, luciferase activity was measured according to the manufacturer’s protocol using a luciferase reporter assay kit (Promega, Madison, WI, USA) with a luminometer (DE/Centro LB960, Berthold Technologies, Oak Ridge, TN, USA).

### Statistics

Statistical analyses were performed using Prism software version 7.0 (GraphPad, La Jolla, CA, USA). Two-way analysis of variance (ANOVA) or Student’s *t*-tests were performed to determine statistical significance. A value of *p* < 0.05 was considered statistically significant. Further experimental details are available in the Supplemental Information.

## Acknowledgments

This work was supported by Basic Science Research Program through the National Research Foundation of Korea (NRF) funded by the Ministry of Science, ICT & Future Planning (NRF-2018M3A9C8020027 to SK and JWL and 2017R1A2B3005015 to JWL) for the Tumor Microenvironment GCRC (2011-0030001) to JWL.

## Supplemental Information

Supplementary table and figures are found at the journal’s web site.

## Author contributions

JR and EK performed most experiments; EK and MKK helped animal experiments; JWJ, SHN, JEK, DGS, and HJK helped with imaging experiments and with reagents; JHL, JHY, and HYK helped with clinical patient tissues; TS helped with the RNA-Seq analysis; SK helped with anti-TM4SF5 antibodies; JWL designed the experiments and wrote the manuscript.

## The paper explained

### PROBLEM

Nonalcoholic liver disease is a chronic disease with immuno-metabolic disorders during NAFLD progression to NASH/fibrosis, cirrhosis and cancer. The molecular mechanisms for the earlier progression to NASH/fibrosis remain unclear.

### RESULTS

*Tm4sf5*-engineered mice revealed that TM4SF5-dependent SIRT1 modulation and chemokines promoted NAFLD, NASH, and fibrosis in an age-dependent manner. TM4SF5/STAT3-mediated laminin expression in hepatocytes critically promoted the earlier progression toward NASH.

### IMPACT

TM4SF5-mediated SIRT1 modulation differentially linked to steatotic SREBPs or NASH-fibrotic STAT3 signaling pathways mechanistically coverts NAFLD toward NASH and fibrosis involving collagen and laminin expressions in HSCs and hepatocytes, respectively.

## References

Agarwal R, Agarwal P (2017) Targeting extracellular matrix remodeling in disease: Could resveratrol be a potential candidate? Exp Biol Med (Maywood) 242: 374–383

Aishima S, Matsuura S, Terashi T, Taguchi K, Shimada M, Maehara Y, Tsuneyoshi M (2004) Aberrant expression of laminin gamma 2 chain and its prognostic significance in intrahepatic cholangiocarcinoma according to growth morphology. Mod Pathol 17: 938–945

Anstee QM, Reeves HL, Kotsiliti E, Govaere O, Heikenwalder M (2019) From NASH to HCC: current concepts and future challenges. Nat Rev Gastroenterol Hepatol 16: 411–428

Ateş M, Alpaslan Pınarlı F, Take Kaplanoğlu G, Tiryaki M, Mercan S, Erdoğan D, Böyük G, Fırat Z, Eyerci N, Topaloğlu O et al (2012) A Modified Method for Isolation of Rat Hepatocyte: Saving Time Increases Viability. Niche 1: 8–11

Biancheri P, Giuffrida P, Docena GH, MacDonald TT, Corazza GR, Di Sabatino A (2014) The role of transforming growth factor (TGF)-beta in modulating the immune response and fibrogenesis in the gut. Cytokine Growth Factor Rev 25: 45–55

Bonnans C, Chou J, Werb Z (2014) Remodelling the extracellular matrix in development and disease. Nat Rev Mol Cell Biol 15: 786–801

Gong Z, Tas E, Yakar S, Muzumdar R (2017) Hepatic lipid metabolism and non-alcoholic fatty liver disease in aging. Mol Cell Endocrinol 455: 115–130

Gressner AM, Weiskirchen R (2006) Modern pathogenetic concepts of liver fibrosis suggest stellate cells and TGF-beta as major players and therapeutic targets. J Cell Mol Med 10: 76–99

Houtkooper RH, Pirinen E, Auwerx J (2012) Sirtuins as regulators of metabolism and healthspan. Nat Rev Mol Cell Biol 13: 225–238

Jung JE, Lee HG, Cho IH, Chung DH, Yoon SH, Yang YM, Lee JW, Choi S, Park JW, Ye SK et al (2005) STAT3 is a potential modulator of HIF-1-mediated VEGF expression in human renal carcinoma cells. Faseb J 19: 1296–1298

Jung JW, Macalino SJY, Cui M, Kim JE, Kim HJ, Song DG, Nam SH, Kim S, Choi S, Lee JW (2019) Transmembrane 4 L Six Family Member 5 Senses Arginine for mTORC1 Signaling. Cell Metab 29: 1306–1319 e1307

Jung O, Choi S, Jang SB, Lee SA, Lim ST, Choi YJ, Kim HJ, Kim DH, Kwak TK, Kim H et al (2012) Tetraspan TM4SF5-dependent direct activation of FAK and metastatic potential of hepatocarcinoma cells. J Cell Sci 125: 5960–5973

Jung O, Choi YJ, Kwak TK, Kang M, Lee MS, Ryu J, Kim HJ, Lee JW (2013) The COOH-terminus of TM4SF5 in hepatoma cell lines regulates c-Src to form invasive protrusions via EGFR Tyr845 phosphorylation. Biochim Biophys Acta 1833: 629–642

Kang M, Choi S, Jeong SJ, Lee SA, Kwak TK, Kim H, Jung O, Lee MS, Ko Y, Ryu J et al (2012a) Cross-talk between TGFbeta1 and EGFR signalling pathways induces TM4SF5 expression and epithelial-mesenchymal transition. Biochem J 443: 691–700

Kang M, Jeong SJ, Park SY, Lee HJ, Kim HJ, Park KH, Ye SK, Kim SH, Lee JW (2012b) Antagonistic regulation of transmembrane 4 L6 family member 5 attenuates fibrotic phenotypes in CCl(4) -treated mice. FEBS J 279: 625–635

Kiyokawa H, Yasuda H, Oikawa R, Okuse C, Matsumoto N, Ikeda H, Watanabe T, Yamamoto H, Itoh F, Otsubo T et al (2017) Serum monomeric laminin-gamma2 as a novel biomarker for hepatocellular carcinoma. Cancer Sci 108: 1432–1439

Lee JW (2015) Transmembrane 4 L Six Family Member 5 (TM4SF5)-Mediated Epithelial-Mesenchymal Transition in Liver Diseases. Int Rev Cell Mol Biol 319: 141–163

Lee SA, Lee SY, Cho IH, Oh MA, Kang ES, Kim YB, Seo WD, Choi S, Nam JO, Tamamori-Adachi M et al (2008) Tetraspanin TM4SF5 mediates loss of contact inhibition through epithelial-mesenchymal transition in human hepatocarcinoma. J Clin Invest 118: 1354–1366

Lee SA, Ryu HW, Kim YM, Choi S, Lee MJ, Kwak TK, Kim HJ, Cho M, Park KH, Lee JW (2009) Blockade of four-transmembrane L6 family member 5 (TM4SF5)-mediated tumorigenicity in hepatocytes by a synthetic chalcone derivative. Hepatology 49: 1316–1325

Li BH, He FP, Yang X, Chen YW, Fan JG (2017) Steatosis induced CCL5 contributes to early-stage liver fibrosis in nonalcoholic fatty liver disease progress. Transl Res 180: 103–117 e104

Maher JJ, McGuire RF (1990) Extracellular matrix gene expression increases preferentially in rat lipocytes and sinusoidal endothelial cells during hepatic fibrosis in vivo. J Clin Invest 86: 1641–1648

Marra F, Tacke F (2014) Roles for chemokines in liver disease. Gastroenterology 147: 577–594 e571

Moon YA (2017) The SCAP/SREBP Pathway: A Mediator of Hepatic Steatosis. Endocrinol Metab (Seoul) 32: 6–10

Roskams T DV, Verslype C. (2007) Development, structure and function of the liver. In MacSween’s Pathology of the Liver. 5th edn., Burt AD PB, Ferrell LD, (ed) pp 1–74. Philidelphia, PA, USA: Churchill Livingstone Elsevier

Ryu J, Kang M, Lee M-S, Kim H-J, Nam SH, Song HE, Lee D, Lee JW (2014) Cross Talk between the TM4SF5/Focal Adhesion Kinase and the Interleukin-6/STAT3 Pathways Promotes Immune Escape of Human Liver Cancer Cells. Mol Cell Biol 34: 2946–2960

Sanyal AJ (2019) Past, present and future perspectives in nonalcoholic fatty liver disease. Nat Rev Gastroenterol Hepatol 16: 377–386

Sircana A, Paschetta E, Saba F, Molinaro F, Musso G (2019) Recent Insight into the Role of Fibrosis in Nonalcoholic Steatohepatitis-Related Hepatocellular Carcinoma. Int J Mol Sci 20

Taura K, Miura K, Iwaisako K, Osterreicher CH, Kodama Y, Penz-Osterreicher M, Brenner DA (2010) Hepatocytes do not undergo epithelial-mesenchymal transition in liver fibrosis in mice. Hepatology 51: 1027–1036

Tu T, Calabro SR, Lee A, Maczurek AE, Budzinska MA, Warner FJ, McLennan SV, Shackel NA (2015) Hepatocytes in liver injury: Victim, bystander, or accomplice in progressive fibrosis? J Gastroenterol Hepatol 30: 1696–1704

Yang MC, Wang CJ, Liao PC, Yen CJ, Shan YS (2014) Hepatic stellate cells secretes type I collagen to trigger epithelial mesenchymal transition of hepatoma cells. Am J Cancer Res 4: 751–763

Zeisberg M, Yang C, Martino M, Duncan MB, Rieder F, Tanjore H, Kalluri R (2007) Fibroblasts derive from hepatocytes in liver fibrosis via epithelial to mesenchymal transition. J Biol Chem 282: 23337–23347

Zhong W, Shen WF, Ning BF, Hu PF, Lin Y, Yue HY, Yin C, Hou JL, Chen YX, Zhang JP et al (2009) Inhibition of extracellular signal-regulated kinase 1 by adenovirus mediated small interfering RNA attenuates hepatic fibrosis in rats. Hepatology 50: 1524–1536

